# Sex chromosome dominance in a UV sexual system

**DOI:** 10.1101/2023.12.28.573518

**Authors:** Jeromine Vigneau, Claudia Martinho, Olivier Godfroy, Min Zheng, Fabian B. Haas, Michael Borg, Susana M. Coelho

## Abstract

The alternation between multicellular haploid gametophytes and diploid sporophytes is a defining feature of most plant and algal life cycles. In such organisms, male and female sexes are determined in the haploid gametophyte with a female (U) or male (V) sex chromosome. Once the U and V chromosomes unite at fertilisation, sex determination no longer occurs, raising key questions about the fate of UV sex chromosomes in the diploid sporophyte stage of the life cycle. Here, we unravel the genetic and molecular interactions between the U and V chromosomes by assessing transcriptional and chromatin states across the life cycle of the brown alga *Ectocarpus* alongside *ouroboros* mutants that decouple life cycle stage from ploidy. We reveal how sex chromosome genes are developmentally regulated across the life cycle, with genes involved in female sex determination in particular undergoing strong down-regulation in the sporophyte. Diploid *ouroboros* mutants containing both a U and V sex chromosome behave as functional male gametophytes yet still exhibit feminized transcription, suggesting that presence of the V chromosome alone is insufficient to fully suppress female developmental program. Although the silencing of sex chromosome genes in the diploid sporophyte does not appear to correlate with localised changes in chromatin state, small RNAs may play a role in the repression of a female sex-linked gene. Finally, we show how histone H3K79me2 is globally re-configured in the diploid phase of the life cycle, including the sex determining region of the UV sex chromosomes. Contrary to its pattern in the haploid gametophyte, H3K79me2 no longer associates with repressed genes in the diploid sporophyte, suggesting that the function of this histone mark in *Ectocarpus* may be more complex than previously appreciated.

## Introduction

The sex of an organism is usually determined either through environmental cues or genetically by means of a sex locus or sex chromosome. A key factor that influences the nature of genetic sex determination is the stage of the life cycle when this occurs. In animals and flowering plants with XX/XY or ZW/ZZ systems, sex is determined during the diploid phase of the life cycle. However, in many other eukaryotes like bryophytes and most brown, red and green algae, sex is determined during a haploid phase of the life cycle called the gametophyte generation (Coelho et al., 2018). The brown algae provide a classic example of a haploid-diploid life cycle with U/V sex determination (Coelho et al., 2007; Coelho et al., 2018). Haploid sex determination in these organisms occurs at meiosis and is dependent on the inheritance of a U or V sex chromosome that will dictate whether the gametophyte develops into a female or a male, respectively (Coelho and Umen, 2021; Coelho et al., 2018). Once fully developed, the gametophytes will undergo gametogenesis to produce gametes via mitosis. Once released into seawater, male gametes are attracted by a pheromone released from the female gamete (Maier, 1995), which then fuse to reconstitute a diploid genome in the sporophyte generation. Diploid sporophytes thus carry both a U and V sex chromosome yet are phenotypically ‘asexual’ since they bear no sexual characteristics but rather undergo meiosis to re-initiate the haploid phase and complete the life cycle.

The brown alga *Ectocarpus* has recently emerged as a powerful model to study the structure and evolution of UV sex chromosomes. The *Ectocarpus* U and V chromosomes are characterized by a distinct sex-determining region (SDR) of similar size and structure that each contains about 20 genes (Ahmed et al., 2014; Coelho et al., 2019). Nine and eight of these genes are sex-specific and are only present on the SDR of the U or V chromosome, respectively. A further 12 gametolog genes are shared by both SDRs, which are remnants of the ancestral pair of autosomes that gave rise to the UV sex chromosomes (Ahmed et al., 2014; Lipinska et al., 2017). While recombination is suppressed within the SDRs, the large majority of the U and V recombine along pseudoautosomal regions (PAR) that flank the centrally-located SDR (Avia et al., 2018; Luthringer et al., 2015b). Compared with autosomes, the PAR tends to harbour an excess of evolutionary young or taxonomically restricted genes while each SDR is highly enriched for repeats and transposable elements (TEs) (Ahmed et al., 2014; Luthringer et al., 2015b). The unique genetic makeup of the PAR and SDR also manifests a highly distinct chromatin landscape compared to autosomes, which is enriched for repressive histone H3K79 and H4K20 methylation in *Ectocarpus* (Gueno et al., 2022).

Theoretical models have long predicted that genes within the SDR would have sex-specific functions and would be expressed specially in the haploid phase of the life cycle (Bull, 1978). Indeed, expression profiling in UV systems like brown algae and moss has revealed a preferential expression of SDR genes in the gametophyte stage (Ahmed et al., 2014; Avia et al., 2018; Carey et al., 2021; Lipinska et al., 2017). Given that phenotypic sex is not expressed in the sporophyte generation, it is unlikely that SDR genes would be required during the diploid phase of the life cycle, although such predictions have yet to be tested experimentally. Indeed, very few studied have addressed the potential genetic interactions between U and V sex-linked genes.

Brown algae are ideal to study the genetic interactions of the U and V chromosomes because ploidy levels are not always correlated with a particular stage in the life cycle. For example, the gametes in several algal species can develop into haploid sporophytes through parthenogenesis (Arun et al., 2019; Bothwell et al., 2010). Conversely, the gametophyte generation is continuously reiterated regardless of ploidy in mutants of *OUROBOROS (ORO)*, which encodes a TALE-HD transcription factor that is required for the gametophyte-to-sporophyte transition in *Ectocarpus* (Arun et al., 2019; Coelho et al., 2011). Strikingly, diploid *oro;oro* mutants bear both a U and V sex chromosome yet phenotypically behave like a male gametophyte, suggesting that the SDR of the male V chromosome is dominant over that of the female U (Ahmed et al., 2014). The apparent dominance of the male SDR is suggestive of a master regulatory gene that triggers male development and/or suppresses female development, with the most likely candidate being a male SDR-specific gene encoding a high-mobility group (HMG) transcription factor (Lipinska et al., 2017; Müller et al., 2021). The observation that the V acts dominantly over the U in a gametophyte context thus provides a key opportunity to gain further insight into the genetic and molecular mechanisms underlying UV sex determination. How are SDR genes and sexual differentiation programs regulated across the life cycle? What is the molecular basis for the male gametophyte-like phenotype of *oro;oro* mutants and what can this tell us about the genetic interactions of the UV chromosomes? And given its pervasive role in XX/XY sex determination systems, does any form of dosage compensation also occur on the U and V sex chromosomes during the diploid sporophyte generation?

Here, we examine the developmental regulation and intricate relationship of the U and V sex chromosomes across the life cycle of *Ectocarpus.* To unravel these genetic interactions, we investigate gene expression and chromatin state in both wild type (WT) and *oro* mutants that uncouple life cycle stage from ploidy. We show how sex chromosome genes are developmentally regulated across the life cycle, with female sex-linked genes on the U chromosome strongly repressed in the diploid phase. We further identify at least one male sex-linked gene that is subjected to dosage compensation in the sporophyte. Despite the homozygous loss of ORO in the diploid phase resulting in a male gametophyte-like phenotype, we show how these mutants exhibit feminized transcriptional patterns, suggesting that ORO activity is required to suppress the expression of sex-linked genes on both the U and V sex chromosome. SDR gene regulation in the sporophyte does not appear to involve local changes in chromatin state but may involve small RNA silencing at one female-specific gene. Finally, we show how histone H3K79me2 modifications are globally re-configured across the *Ectocarpus* life cycle, with drastic reductions observed over the SDR of the U and V sex chromosomes. Contrary to its pattern in the gametophyte generation, H3K79me2 deposition does not correlate with repressed genes in the sporophyte, suggesting a distinct role for this histone mark in the diploid phase of the life cycle.

## Results

### ORO is required to suppress male sex determination in the diploid phase of the life cycle

To investigate interactions between the U and V sex chromosomes, we exploited *oro* mutant lines in *Ectocarpus* that uncouple life cycle stage from ploidy (Fig. 1A) (Coelho et al., 2011). We have shown previously how haploid male and female *oro* gametophytes largely phenocopy wild type (WT) gametophytes at a morphological and functional level (Fig. 1B) (Arun et al., 2019; Coelho et al., 2011). To explore the effect of *oro* mutations on gametophytic gene expression, we compared gene expression of *oro* mutants with WT gametophyte lines (Fig. 1C-D; Table S1). We observed a large number of differentially expressed genes (DEGs) in *oro* males and *oro* females compared with WT gametophytes of the equivalent sex (Fig. 1D; Table S1). Around a half of these ORO-dependent DEGs were common to both sexes (46.0% in males; 48.6% in female) (Fig. 1E), suggesting that many of the transcriptional changes observed in the absence of ORO largely occur independent of sexual differentiation. Consistently, the sex bias of effector genes involved in specifying a WT sexual phenotype (i.e., sex-biased genes) remain sex-biased in *oro* mutants (Fig. 1F), indicating that *oro* gametophytes retain male and female characteristics at a transcriptional level. Taken together, these results confirmed that the *oro* mutation does not affect the sexual identity of haploid gametophytes.

**Figure 1.**
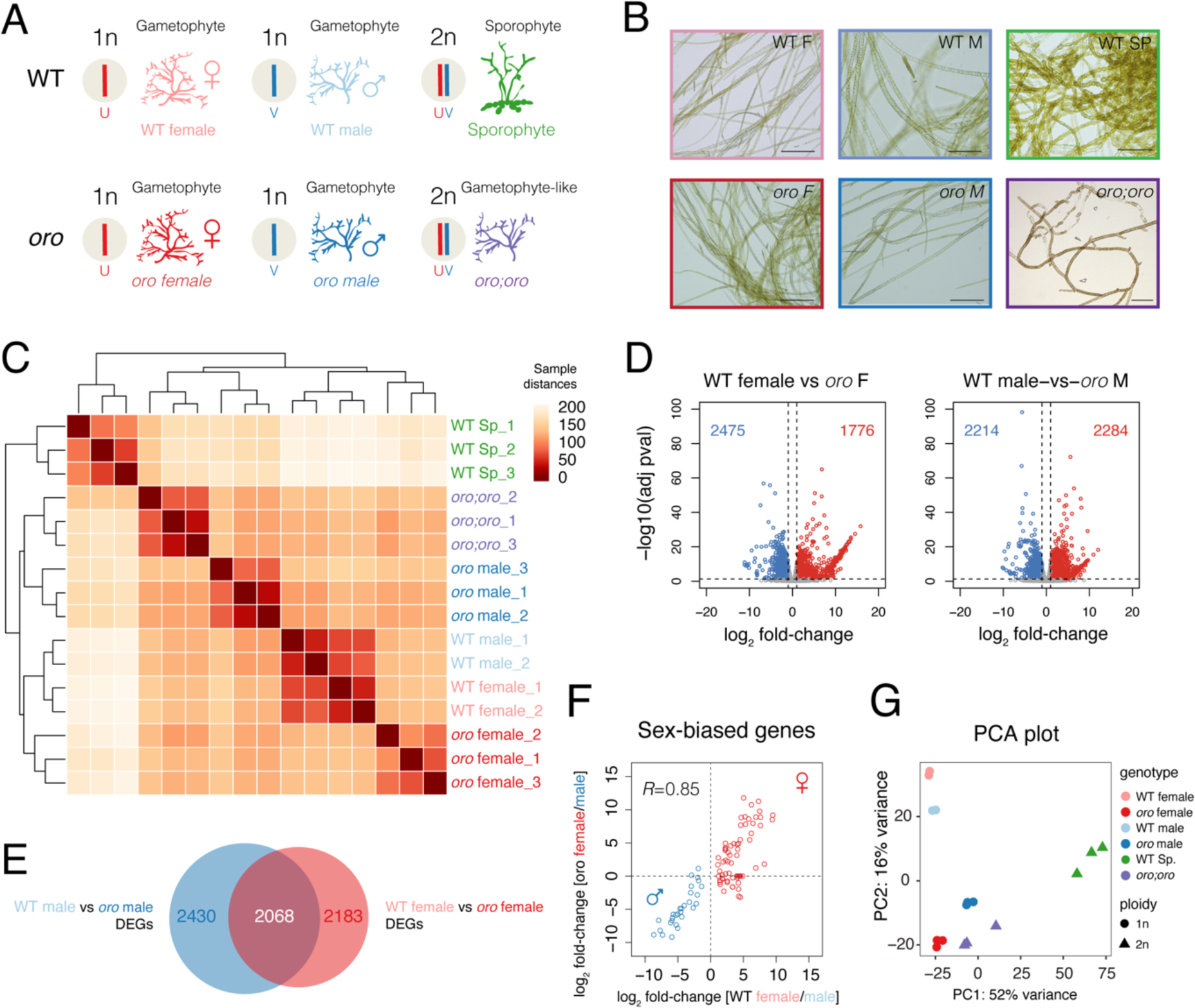
Transcriptional profiling across the *Ectocarpus* life cycle. (A) Schematic diagram summarising the genotypes profiled in this study. (B) Representative images of each genotype illustrate the gametophyte-like phenotype of diploid *oro;oro* mutants. Scale bars = 100μm. (C) Sample distance matrix of the RNA-seq datasets generated and analysed in this study. (D) Volcano plot summarising differentially-expressed genes (DEGs) between WT and *oro* female (left) and male (right) gametophytes. Up-regulated (log2 fold-change > 1 and adjusted *P* value < 0.05) and down-regulated (log2 fold-change < −1 and adjusted *P* value < 0.05) genes are highlighted in red and blue, respectively. The total number of DEGs are indicated in each plot. (E) Venn diagram summarising the overlap of DEGs between WT and *oro* female and male gametophytes. (F). Plot of the pairwise correlation of the differential expression (log2 fold-change) of sex-biased genes (SBGs) between female and males in a WT and *oro* background. Female and male SBGs are highlighted in red and blue, respectively. Pearson’s correlation coefficient is shown. (G) Principal component analysis illustrating the variation among the RNA-seq datasets from WT females (n = 2 replicates) (Gueno et al., 2022), WT males (n = 2 replicates) (Gueno et al., 2022), WT sporophytes (n = 3 replicates), haploid *oro* females (n = 3 replicates), haploid *oro* males (n = 3 replicates) and diploid *oro;oro* mutants (n = 3 replicates).

Male and female *oro* gametophytes were grown to fertility and crossed to obtain a diploid (UV) *oro;oro* mutant line (line Ec581) (Ahmed et al., 2014). This allowed us to combine the U and V chromosomes within the same gamete-producing individuals (Fig. 1A). Consistent with the continuous iteration of gametophytes in *oro* lines via parthenogenesis (Coelho et al., 2011), diploid *oro;oro* mutants also retained gametophyte-like characteristics including richly-branched wavy thalli and a lack of substrate adherence (Fig. 1B). This was also reflected at the level of gene expression, with diploid *oro;oro* mutants more closely related to haploid gametophytes than to WT diploid sporophytes (Fig. 1C,G). Test crosses using the diploid *oro;oro* line crossed with female lines suggested that it behaves as a functional male gametophyte (Fig. S1), confirming previous results (Ahmed et al., 2014). Thus, the homozygous loss of ORO in diploid lines results in functional gametophytes that phenocopy male sex characteristics, despite the presence of both the U and V sex chromosome.

### Developmental transcription factors undergo dynamic regulation across the life cycle

The phenotype observed in diploid *oro;oro* gametophyte-like mutants indicates that the V chromosome is genetically dominant and leads to a male sexual phenotype. This suggests that the V chromosome harbours a dominant genetic factor that is responsible to specify male gametophytic fate. Consistent with this idea, our recent work has revealed that a gene present on the V chromosome encodes a high-mobility group (HMG) transcription factor called HMG-sex that acts a master regulator of male sex determination in *Ectocarpus* and kelps (Luthringer et al, submitted). Consistently, *HMG-sex* transcripts were barely detectable in WT sporophytes compared to male gametophytes (Fig. 2A-C). Interestingly, while *HMG-sex* expression was reduced in the WT sporophyte, it was increased two-fold in both haploid and diploid *oro* mutants compared with the WT male gametophyte (Fig. 2A). This indicates that ORO activity is required to suppress *HMG-sex* expression in the WT sporophyte in a manner similar to its neighbouring genes on the male SDR (Fig. 2B-C). Given the role *HMG-sex* plays in male sex determination (Luthringer et al., submitted), its increased expression in diploid *oro;oro* mutants likely explains the male sex phenotype observed in these lines.

**Figure 2.**
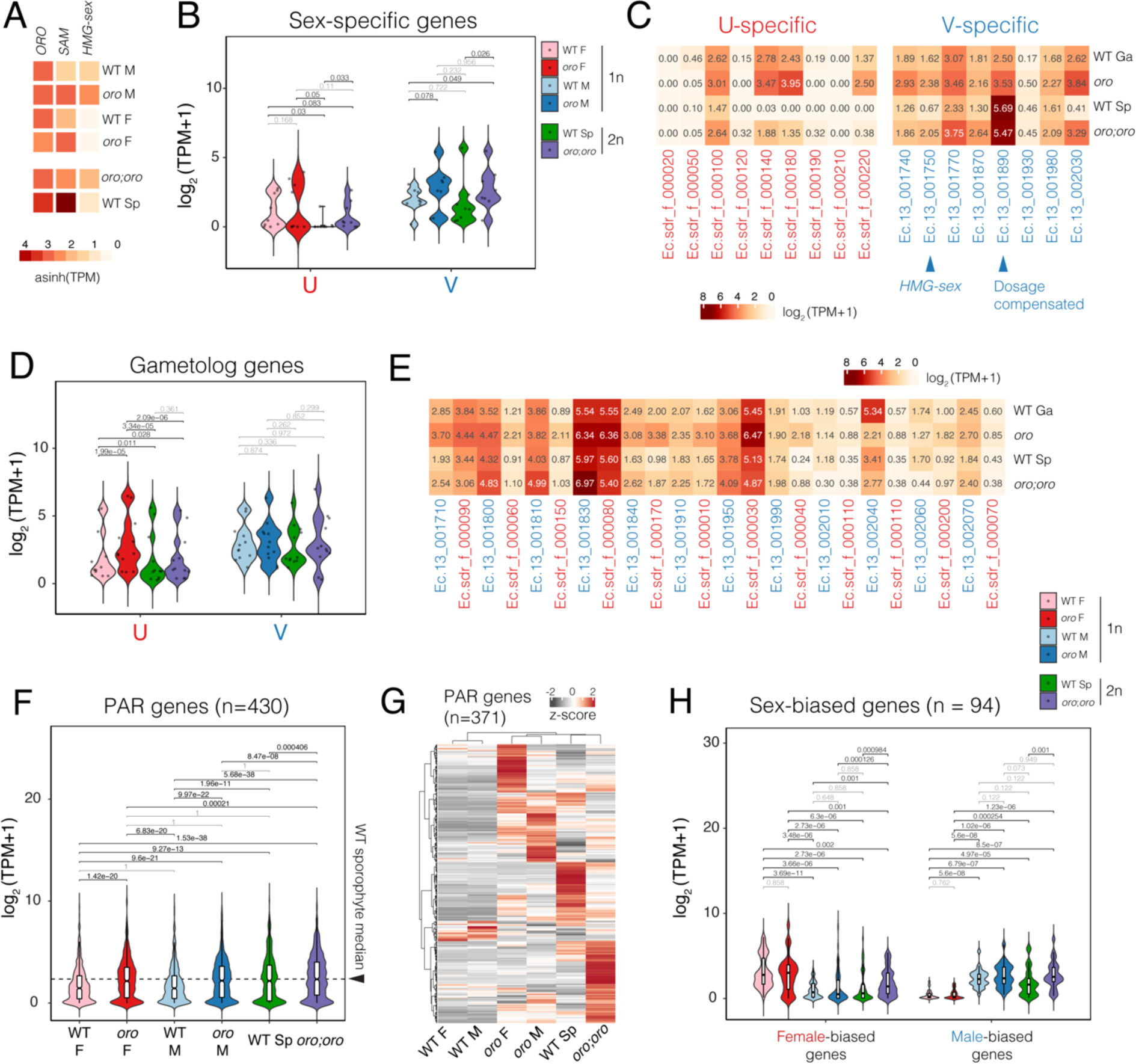
Transcriptional dynamics of the sex chromosomes across the *Ectocarpus* life cycle. (A) Expression of the transcription factors genes *ORO*, *SAM* and *HMG-sex* across the WT and *oro* genotypes and life stages. Expression represents the inverse hyperbolic sine (asinh) transform of the mean RNA-seq TPM values. The mean value of the biological replicates is shown. (B-E) Violin plots and heatmaps summarising the expression of (B-C) genes specific to the sex-determining region (SDR) of the female U and male V sex chromosomes and (D-E) gametolog genes. Expression represents the log2 of the mean RNA-seq TPM+1 values. (F-G) Violin plot and heatmap summarising the expression of genes located on the pseudoautosomal (PAR) region. Expression in the violin plot represents the log2 of the mean RNA-seq TPM+1 values, while the heatmap represents z-score normalised RNA-seq TPM values. (H) Violin plot summarising the expression of sex-biased genes (n = 94). Expression represents the log2 of the mean RNA-seq TPM+1 values. Statistical pairwise comparisons indicated on the violin plots represent the *P* value of paired student t-tests (B,D,F) or paired Wilcoxon signed-rank tests (H). Significant (*P* < 0.05) and non-significant (*P* > 0.05) comparisons are indicated in black and grey, respectively.

While *ORO* expression is fairly stable across major stages of the *Ectocarpus* life cycle, *SAMSARA* (*SAM*) has a complementary expression pattern when compared with *HMG-sex* (Fig. 2A). *SAM* encodes a TALE-HD transcription factor that is also required for the gametophyte-to-sporophyte transition in *Ectocarpus* (Arun et al., 2019). The similar phenotype of *oro* and *sam* mutants, coupled with the fact that ORO and SAM can interact *in vitro* (Arun et al., 2019), suggests that they form a heterodimer to exert their function in a manner similar to other TALE-HD transcription factors (Dierschke et al., 2021). Both *ORO* and *SAM* had the highest expression level in the WT sporophyte generation, which also happens to coincide with downregulation of *HMG-sex* (Fig. 2A), suggesting that ORO/SAM activity is associated to repression of *HMG-sex* expression, be this directly or indirectly.

### UV sex chromosome genes are developmentally regulated across the *Ectocarpus* life cycle

Genes that function to specify a female or male gametophyte are predicted to be retained within the SDR of the U or V sex chromosomes, respectively. In contrast, sporophyte-specific genes required in the diploid phase would be lost from one of the SDRs if hemizygosity were to not greatly reduce fitness (Bull, 1978). To evaluate these predictions and further understand the mechanism underlying dominance of the V over the U chromosome, we examined the patterns of sex chromosome gene expression across several stages of the *Ectocarpus* life cycle.

In WT sporophytes, where no sex is determined, transcript abundance of both U- and V-specific genes (i.e., genes that are present specifically on the female or male SDR, respectively) was reduced compared to WT gametophytes, with U-specific genes being largely silenced (Fig. 2B-C). Only one of the nine U-specific genes remained appreciably expressed (TPM >1) in WT sporophytes, compared to five of the eight V-specific genes (Fig. 2C). In contrast, V-specific genes were significantly upregulated in haploid *oro* mutants compared to WT male gametophytes (Fig. 2B-C). Interestingly, the expression of both U- and V-specific genes was significantly increased in diploid *oro;oro* mutants compared to WT sporophytes, with V-specific genes regaining transcript levels similar to those in WT male gametophytes (Fig. 2B-C). ORO activity appears thus to be required to suppress SDR gene expression during the diploid phase of the life cycle, further explaining the gametophyte-like identity of *oro;oro* mutants.

Because dosage compensation is crucial to equalise sex chromosome gene expression in organisms with XX/XY sex determination (Disteche, 2012), we wondered whether this would also be the case in a UV sex determination system. Although chromosome-level dosage compensation of the U and V would not be required during the haploid phase of the life cycle, gene-level dosage compensation could take place for genes that are specific to either the U or V in the diploid sporophyte generation. Closer examination of the gene located on the SDR of each sex chromosome revealed only one dosage-compensated gene among the male SDR (*Ec-13_001890*, Fig. 2C). This locus encodes a hypothetical protein with a transmembrane domain and had double the expression level in the sporophyte compared to the gametophyte (Fig. 2C). No dosage compensated genes were identified among U-specific genes. Thus, dosage compensation of SDR genes appears to be rare in the diploid phase of the *Ectocarpus* life cycle.

We next examined the expression of gametolog genes, which represent a class of homologous gene pairs present on the SDR of both the U and V chromosomes (Ahmed et al., 2014). Interestingly, gametologs on the U chromosome were significantly down-regulated in WT sporophytes compared with WT female gametophytes (Fig. 2D-E). In contrast, no significant changes in transcript abundance were observed across the life cycle at V gametologs (Fig. 2D-E). Moreover, unlike U-specific genes, U gametologs were not significantly up-regulated in diploid *oro;oro* mutants compared with WT sporophytes (Fig. 2D-E). This observation suggests that the presence of both sex chromosomes in the sporophyte is associated with the silencing of some U (but not V) gametologs. Together, our results indicate that genes located within the SDR of both the U and V chromosome are predominantly expressed in the gametophyte generation, with those on the U most strongly repressed in the diploid phase.

We extended our analysis to genes present within the PARs of the sex chromosomes, which are identical in both the U and V chromosomes (Avia et al., 2018; Luthringer et al., 2015b). Consistent with our previous findings (Luthringer et al., 2015b), PAR genes had higher levels of expression in WT sporophytes compared with WT male or female gametophytes (Fig. 2F). Interestingly, PAR transcripts were significantly upregulated in *oro* male or female gametophytes compared with their WT equivalent (Fig. 2F), suggesting that ORO may be required to suppress PAR gene expression in the haploid phase of the life cycle. PAR gene expression was also modestly but significantly increased in diploid *oro;oro* mutants compared to WT sporophytes (Fig. 2F), indicating that ORO may also modulate PAR gene expression in the diploid phase of the life cycle albeit to a lesser degree than in the haploid phase. Indeed, closer examination of PAR gene expression revealed that the majority (86.2%; 371 of 430) are subjected to substantial ORO-dependent repression in both haploid and diploid phases (Fig. 2G). Thus, unlike the attenuated expression of SDR genes in the diploid phase, PAR genes appear to be downregulated during the haploid phase of the life cycle in an ORO-dependent manner.

### Autosomal sex-biased genes are repressed in the sporophyte generation

We further assessed the impact of increased ploidy on the expression of sex-biased genes (SBGs), which are defined as autosomal genes that are differentially regulated between male and female gametophytes. We considered a conservative set of 94 SBGs that were sex-biased in at least two out of three three independent datasets (Table S2) (Gueno et al., 2022; Lipinska et al., 2015). SBGs are unlikely to be required in the sporophyte generation given the lack of expression of sexual phenotypes during this stage of the life cycle (Luthringer et al., 2015a). Indeed, both male and female SBGs were significantly down-regulated in WT sporophytes compared to gametophytes of the equivalent sex (Fig. 2H). As was observed for genes specific to the SDR of the female U chromosome (Fig. 2B), the strong reduction in transcript levels in WT sporophytes was more pronounced for female SBGs compared to male SBGs (Fig. 2H). Interestingly, male SBG expression was once more increased in diploid *oro;oro* mutants compared to WT sporophytes, reaching levels similar to those in the WT male gametophyte (Fig. 2H). Thus, the expression of V-specific genes and male SBGs in diploid *oro;oro* gametophytes resembles that in haploid male gametophytes, likely explaining the ability of *oro;oro* mutants to function as fertile male (Fig. S1).

Despite diploid *oro;oro* mutants functioning as male gametophytes, diploid *oro;oro* mutants also appear to be ‘feminized’ by the increased expression of a subset of female SBGs (Fig. 2H). This indicates complex interplay between the differentiation programs governing each sex, despite the male program being functionally and genetically dominant over that of the female, which is likely reflective of the presence of the U chromosome in diploid gametophytes. In contrast, no significant differences in SBG expression were observed between WT and *oro* haploid gametophytes (Fig. 2H). Thus, consistent with the presence of both a U and V sex chromosome, *oro;oro* gametophytes phenocopy male sex but molecularly display a slightly feminised transcriptome (Fig. 2H). These results further emphasise how ORO activity is required to suppress gametophytic fate in the diploid phase of the life cycle via the repression of both sex-specific and sex-biased genes.

### SDR gene regulation is largely independent of sRNA silencing and local chromatin changes

Small RNAs (sRNAs) play an important role in the regulation of gene expression and TE silencing in eukaryotes (Chen, 2009; Rana, 2007). We therefore investigated whether the differential expression of SDR genes between WT sporophytes and *oro;oro* diploid mutants was associated with differences in sRNA accumulation (Table S3). Overall, sRNAs do not appear to explain gene expression changes at the female SDR (Fig. S2A-B). Conversely, global sRNA accumulation of multi mapped sRNA was moderately but significantly decreased in *oro;oro* compared to the WT sporophyte at the male SDR (Fig. S2B). Accordingly, one-third (8/23; 34.8%) of the sRNAs associated with genes on the male SDR were significantly downregulated (*P* < 0.05 and log_2_ fold-change < −1.2) in *oro;oro* mutants compared to the WT sporophyte (Table S3). Although six of these eight genes encoded gametologs (i.e., genes with a homolog on the female SDR), there was no significant enrichment for gametologs among down-regulated genes (Chi-square test *P* = 0.3537, Table S3). To further assess whether sRNAs were negative regulators of SDR gene expression, we calculated the correlation coefficient of the fold-change difference in sRNA accumulation and mRNA abundance between WT sporophytes and *oro;oro* mutants (Fig. 3A). Surprisingly, sRNA accumulation was positively correlated with male SDR gene expression (Fig. 3A), suggesting that sRNAs may not be required for repression of male SDR genes in WT sporophytes (Fig. 3A). In contrast, one female SDR gene (*Ec-sdr_f_000040*) that was specifically upregulated in *oro;oro* happened to be associated with decreased accumulation of sRNAs (Fig. 3B). These sRNA accumulated to a higher extent in WT sporophytes within the last intron of the locus and overlapped with an unclassified repeat (Fig. 3B). Thus, aside for one U-specific gene within the female SDR gene, our observations indicate no clear association between the presence of sRNAs and the silencing of SDR genes.

**Figure 3.**
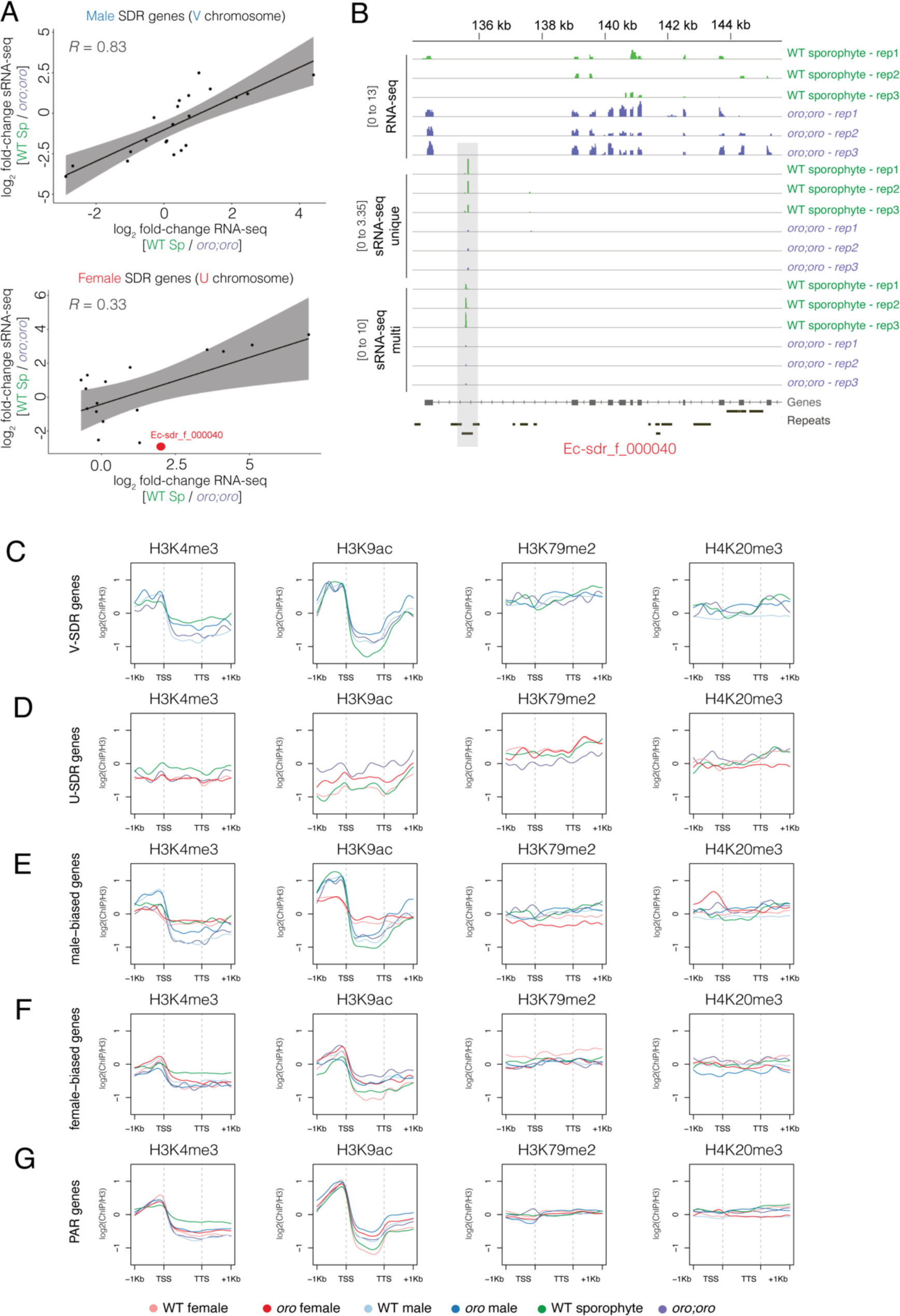
The small RNA and chromatin state of genes involved in sex determination. (A) Plot of the pairwise correlation of the differential expression (log2 fold-change) of male (top) and female (bottom) SDR genes and their associated small RNAs between diploid WT sporophytes and *oro;oro* mutants. Spearmans’s correlation coefficient is shown. Highlighted in red is the female-specific gene *Ec-sdr_f_000040*, which is significantly upregulated in *oro;oro* (log2 fold-change = 1.98; *P =* 0.005) and has a corresponding significant decrease in small RNA levels (log2 fold-change = −2.89; *P =* 4.08×10^-11^). (B) A genome browser view of the *Ec-sdr_f_000040* locus showing RNA-seq coverage alongside unique and multi-mapped small RNAs in diploid WT sporophytes and *oro;oro* mutants. Dynamic small RNAs are highlighted with grey shading. (C-G) The ChIP-seq signal of H3K4me3, H3K9ac, H3K79me2 and H4K20me3 over SDR genes of the female U sex chromosome (C), SDR genes of the male V sex chromosome (D), female-biased genes (E), male-biased genes (F), and PAR genes (G). ChIP-seq signals represent the log2 ratio of immunoprecipitated DNA relative to histone H3 and are colour-coded based on the key at the bottom of Figure 3 to differentiate between the six genotypes profiled in the study.

Recent studies have shown how the re-configuration of histone modifications during sexual differentiation are intimately linked with the establishment and/or maintenance of sex-specific transcriptional programs (Brown and Bachtrog, 2014; Gueno et al., 2022). We thus examined whether the differential deposition of histone modifications was also associated with transcriptional changes of sexual differentiation programs among the different life cycle contexts. Our previous work in *Ectocarpus* has shown that H3K4me3 and H3K9ac are associated with the transcription start sites (TSS) of active genes, while H3K79me2 and H4K20me3 deposition is correlated with decreased transcript abundance (Bourdareau et al., 2021; Gueno et al., 2022). We thus generated ChIP-seq profiles for H3K4me3, H3K9ac, H3K79me2 and H4K20me3 in the four different life cycle stages and compared these with our previously published profiles of WT gametophytes (Fig. S3). We validated the robustness of our ChIP-seq data in two ways. Firstly, we confirmed high correlation between the replicates of each histone mark in each life cycle stage (Fig. S3A-F). Second, we confirmed that the deposition of active and repressive marks was positively and negatively correlated with gene expression, respectively (Fig. S3G-L).

Next, we aggregated ChIP-seq coverage for each histone mark over the different groups of genes involved in sex determination i.e., over U- and V-specific genes and SBGs (Fig. 3C-G). Surprisingly, no major differences in histone mark deposition were evident that could explain the transcriptional changes we observed among the different life cycle contexts for genes located both within the male and female SDR and along the PAR (Fig. 2D-G; Fig. 3C,D,G). However, the upstream promoter region of male SBGs clearly showed reduced levels of active H3K4me3 and H3K9ac marks in WT and *oro* female gametophytes compared with their male equivalent, consistent with the reduced expression of male SBGs in females (Fig. 2H; Fig. 3E). Similarly, the upstream promoter region of female SBGs also showed reduced levels of H3K4me3 and H3K9ac in WT male gametophytes and WT sporophytes (Fig. 3F), consistent with the reduced expression of female SBGs in these stages (Fig. 2H). Thus, while some sex-biased genes appear to undergo dynamic changes in the level of active histone marks between male and female gametophytes, consistent with our previous findings (Gueno et al., 2022), local *cis* changes at chromatin do not appear to coincide with changes in the expression of U and V sex chromosomes genes.

### The landscape of H3K79me2 is globally re-configured across the *Ectocarpus* life cycle

We next considered whether more global changes in chromatin could explain the silencing of SDR genes in the WT sporophyte generation. Our previous work showed that the repeat-rich sex chromosome in *Ectocarpus* is highly enriched for chromatin signatures defined by repression-associated H3K79me2 marks (Gueno et al., 2022). Consistently, we observed very broad domains of H3K79me2 enrichment along the U and V sex chromosomes, with the majority of the SDRs being covered by H3K79me2 in the WT female and male gametophyte (Fig. 4A-B). In the WT sporophyte, these broad domains of H3K79me2 were reduced in intensity compared with the gametophyte generations, with the female SDR showing a particularly pronounced reduction in H3K79me2 levels (Fig. 4A-B). Differential analysis of H3K79me2 peak enrichment confirmed this, with the majority (17/22; 77.3%) of U-specific genes associated with one or more peaks that were significantly depleted in H3K79me2 levels in the female gametophyte compared with the WT sporophyte (Fig. 4C; Table S4). Only one V-specific gene showed this association (Fig. 4D; Table S5), consistent with a more modest decrease in H3K79me2 along the male SDR (Fig. 4B).

**Figure 4.**
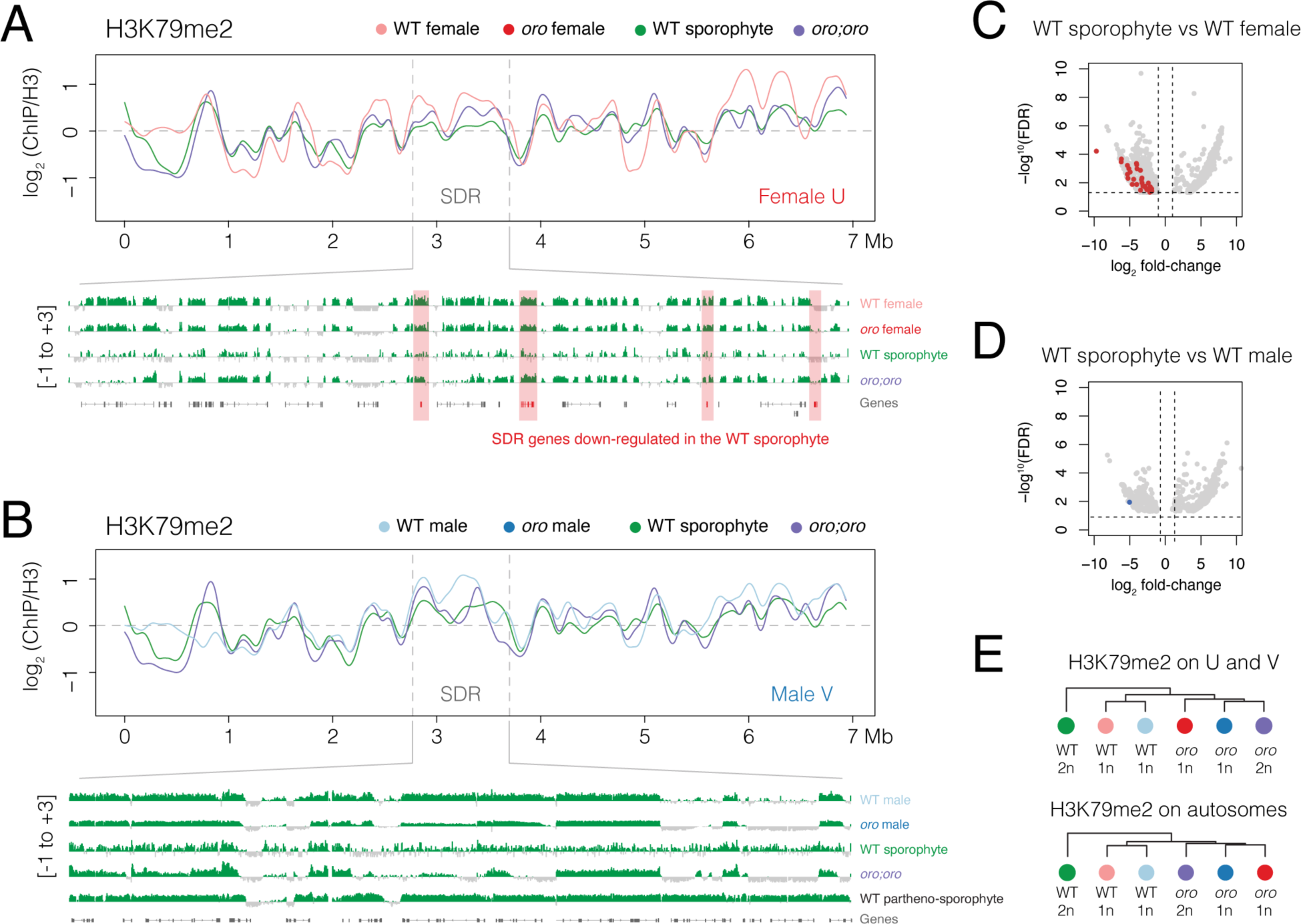
H3K79me2 is globally reconfigured across the *Ectocarpus* life cycle. (A-B) Distribution of H3K79me2 over the female U (A) and male V (B) sex chromosome. The position of the sex-determining region (SDR) of each sex chromosome is indicated between the grey dashed lines. A genome browser view of H3K79me2 enrichment over the complete SDR is provided below. Signals represent the ChIP-seq log2 enrichment of immunoprecipitated H3K79me2-associated DNA relative to histone H3 calculated in 10kb bins. (C-D) Volcano plot of differential enrichment of H3K79me2 peaks between the WT sporophyte and WT female gametophyte (C) and the WT sporophyte and WT male gametophyte (D). Peaks associated with female and male SDR genes are highlighted in red and blue, respectively. (E) Hierarchical clustering of H3K79me2 profiles over sex chromosomes and autosomes.

In contrast, the strong reduction of H3K79me2 within the SDR was much less pronounced in diploid *oro;oro* mutants, which instead more strongly resembled that seen in WT male and female gametophytes (Fig. 4A-B). Similarly, this reduction was not evident in the SDR of a haploid male partheno-sporophyte (Fig. 4B) (Bourdareau et al., 2021), suggesting that the chromosomal reconfiguration of H3K79me2 only occurs in the diploid sporophyte, in the presence of the U chromosome. Hierarchical clustering of H3K79me2 profiles on the U and V sex chromosomes further illustrates how WT sporophytes are clearly distinct from the other life cycle stages, which also manifests along autosomes (Fig. 4E). Thus, the union of the U and V sex chromosome in the sporophyte generation results in a reduction of H3K79me2 along the SDRs, which is once again increased in the gametophyte-like context of diploid *oro;oro* mutants. The global reconfiguration of H3K79me2 is further accompanied by the downregulation of SDR genes in the WT sporophyte generation (Fig. 2B-D), suggesting that this could impact the transcriptional state of the U and V sex chromosome. Closer inspection of H3K79me2 deposition on genes grouped by expression level in the WT sporophyte illustrated a poorer correlation with repressed genes when compared to other life stages (Fig. S3F), further suggesting that the repressive role of H3K79me2 might not extend to the diploid phase of the life cycle.

## Discussion

Eukaryotic organisms that are characterized by an alternation of generations must undergo dynamics changes in genomic activity to specify distinct developmental programs in the alternating haploid and diploid stages of their life cycle (Coelho et al., 2007). As a haploid gametophyte, an individual will grow vegetatively through haploid mitosis and eventually initiate gametogenesis to mate with the opposite sex. A diploid sporophyte will also grow vegetatively through diploid mitoses but will instead initiate meiosis to begin the process of sporulation. Thus, only diploid sporophytes should activate the meiotic program, while haploid gametophytes alone should transition towards gametogenesis. Tight control of these processes must thus be ensured to trigger the correct developmental program at the opportune moment in the life history (Perrin, 2012). Here, we have examined the developmental regulation and intricate relationship of U and V sex chromosomes across the life cycle of the model brown alga *Ectocarpus*.

TALE HD transcription factors are involved in controlling the diploid transition across eukaryotic lineages (Arun et al., 2019; Dierschke et al., 2021; Kariyawasam et al., 2019), whereas sex chromosome genes are predicted to regulate haploid gametophyte development (Bachtrog, 2006; Charlesworth, 2016; Coelho and Umen, 2021). Consistent with this notion, genes involved in haploid to diploid transitions and sex determination, including those on the sex chromosomes, undergo extensive transcriptional reprogramming across the *Ectocarpus* life cycle. This included major developmental regulators like the TALE HD transcription factors ORO and SAM (Arun et al., 2019; Coelho et al., 2011) and the HMG-domain transcription factor HMG-sex.

In male gametophytes, FBG expression remained unchanged compared to the WT sporophyte, suggesting that the male program appears to be driven solely by upregulation of MBGs. In contrast, female gametophytes appear to both strongly downregulate MBGs while strongly upregulating FBGs. Thus, SBGs undergo dynamic and opposing changes in expression during female sexual development, whereas an increase in MBG expression alone appears to be sufficient for male sexual development. This trend is consistent with the male sex determination program acting dominantly over the female program. Moreover, the persistent expression of some FBGs in males suggests that they might also be involved and/or required for aspects of male development. This potential pleiotropic effect of FBGs is further reflected by the fact that FBGs have a broader expression pattern than MBGs in *Ectocarpus* (Cossard et al., 2022; Lipinska et al., 2015). Stronger pleiotropic constraints on the broad expression of FBGs has been shown to be prevalent even among animals models like *Drosophila* (Allen et al., 2018). Our results are consistent with the female sex determination acting as the ‘background’ program of sexual development in *Ectocarpus*, which is then suppressed through the expression of male-dominant regulatory factors (Ahmed et al., 2014; Avia et al., 2018; Müller, 1967; Müller et al., 2021).

Because the U and V sex chromosomes function during the haploid phase of the life cycle, chromosome-scale dosage compensation is not expected to occur as it would in XX/XY sex determination systems (Duan and Larschan, 2019). Nevertheless, we identified one V-specific gene encoding a hypothetical transmembrane protein that appears to be subjected to dosage compensation in the sporophyte generation, suggesting that it might also play an important role in both haploid and diploid phases of the life cycle.

The diploid homozygous loss of the TALE-HD transcription factor ORO results in a male gametophyte-like phenotype that produces functional male gametes capable of undergoing fertilisation, despite the presence of both the U and V sex chromosome (Ahmed et al., 2014). The V chromosome is therefore dominant over the U, which is coherent with the presence of *HMG-sex*, a male sex determining gene present on the SDR of the V chromosome (Luthringer et al. submitted). Interestingly, we observed that *HMG-sex* expression is derepressed in *oro;oro* mutants, suggesting that the regulatory activity of ORO is required to suppress *HMG-sex* expression, be this directly or indirectly. Moreover, diploid *oro;oro* mutants do not fully silence the female developmental program since these lines are transcriptionally “feminized” in comparison to a WT male gametophyte. This is presumably also due to the inability to silence SDR genes in the absence of ORO activity, which would normally occur in the diploid sporophyte generation. Our work thus extends our understanding of how ORO suppresses gametophyte identify in the diploid phase of the *Ectocarpus* life cycle, which could potentially occur via the direct repression of sex determining genes. Future *in vivo* binding studies will hope to further our molecular understanding of how ORO exerts its regulatory role over the U and V sex chromosome.

Our work has further sought to clarify how sexual differentiation programs are differentially regulated across the *Ectocarpus* life cycle. Altogether, our data suggest that the regulation of SDR genes is largely independent of sRNAs since we found no global association between the presence of sRNAs and transcriptional silencing on SDR genes in the sporophyte generation. One notable exception was the U-specific gene *Ec-sdr_f_00040*, which was upregulated in *oro;oro* mutants and was correspondingly correlated to reduced sRNA levels. *Ec-sdr_f_000040* belongs to a highly divergent gametolog pair representing a casein kinase gene that is consistently female-sex linked across a range of brown algal species (Lipinska et al., 2017). Casein kinases are involved in regulating a wide range of developmental processes, including reproduction (Guo et al., 2023; Ogiso et al., 2010; Phadnis et al., 2015; Qu et al., 2021; Zhang et al., 2022), while its orthologue in kelps appears to have an important function in female development (Müller et al., 2021). We speculate that this is gene could also be involved in the transcriptional feminisation of *oro;oro* mutants. Future examination will be required to further understand the intriguing accumulation of sRNAs over the repeat element within the intron of this sex-linked gene.

Despite the transcriptional changes we observed at genes in involved in sex determination, we were puzzled not to find concomitant changes in chromatin state. The transition to sexual development in *Ectocarpus* is characterized by the formation of plurilocular sporangia that produce and release motile gametes (Charrier et al., 2008; Coelho et al., 2007). SDR genes as well as other sex-biased genes are thus only likely to be transcribed within a small subset of reproductive cells and/or during a limited time window of plurilocular sporangia development. Given that our profiles were generated using whole thalli, we speculate whether the excess of vegetative tissue impedes our ability to observe cell-type specific changes in chromatin state. Generating such data is challenging due to the microscopic nature of these reproductive structures in *Ectocarpus*. Future analyses at the single-cell level will hope to clarify how chromatin reprogramming facilitates sexual differentiation during the transition to the haploid gametophyte generation.

Nevertheless, our chromatin profiling has revealed how H3K79me2 deposition is reconfigured across the *Ectocarpus* life cycle. In particular, we observed a strong reduction of H3K79me2 along the SDRs in the sporophyte generation, particularly along the female SDR of the U chromosome, which strongly contrasts with its gametophytic pattern (Bourdareau et al., 2021; Farooq et al., 2016; Gueno et al., 2022). H3K79me2 has been extensively studied in yeast and mammals where it accumulates over the transcribed region of active genes (Nguyen and Zhang, 2011). In contrast, H3K79me2 has an opposing pattern in *Ectocarpus* gametophytes since it preferentially accumulates over transposons and genes with low levels of gene expression (Bourdareau et al., 2021; Gueno et al., 2022). Interestingly, we show here that this association is less evident in the sporophyte since its accumulation is no longer associated with obvious changes in gene expression. This suggests that the role of H3K79me2 may be distinct during the diploid phase of the life cycle and might no longer mediate a repressive role. The global reconfiguration of H3K79me2 during the haploid-diploid transition in *Ectocarpus* is reminiscent of the extensive epigenetic reprogramming observed during haploid-diploid transitions in bryophytes and flowering plants (Vigneau and Borg, 2021), which play a key role in activating pollen-specific genes (Borg et al., 2021; Khouider et al., 2021), sperm specification (Borg et al., 2021) as well as specification of the female gametophyte in *Arabidopsis* (Baroux and Autran, 2015; She and Baroux, 2015; She et al., 2013) Such global chromatin reprogramming events may thus be a general feature of organisms that alternate between haploid and diploid multicellular phases, although future studies are needed to clarify the precise role of H3K79me2 in the brown algal lineage.

## Materials & Methods

### Biological material

The pedigree of the strains used in this study were described previously (Coelho et al. 2011) and correspond to WT male (Ec32) and female (Ec25) gametophytes, WT sporophytes (Ec17), *oro* mutant males (Ec561) and females (Ec560), and diploid *oro;oro* mutants (Ec581). About 10 algal individuals were cultivated in Petri dishes at 14 °C in Provasoli-enriched seawater (Coelho et al., 2012) under a short day regime (12 hours dark / 12 hours light) with 20 μmol photons m^−2^ s^−1^ irradiance.

### RNA-seq analysis

RNA-seq data was generated from culture with same conditions to ensure that the histone PTM and gene expression data were fully compatible. All RNAseq datasets generated in this study were carried out in biological triplicate for each genotype. For each replicate, 10 mg of *Ectocarpus* tissue (∼10 individuals) was patted dry with absorbent tissue paper and flash-frozen. Total RNA was isolated as described previously with minor modifications (Cossard et al., 2022; Lipinska et al., 2015; Wang and Stegemann, 2010). In brief, the tissue was ground to a fine powder in liquid nitrogen in 1.5mL Eppendorf tubes using a micropestle. The algae powder was further homogenized in 700 μL of pre-warmed (at 65°C) Cetlytrimethylammonium Bromide (CTAB)-based extraction buffer (100 mM Tris-HCl pH 8, 1.4 M NaCl, 20 mM EDTA pH 8, 2 % Plant RNA isolation Aid (PVP, Invitrogen #AM9690), 2 % CTAB and 1 % beta-mercaptoethanol) by vortexing and incubated at 65°C until all samples were processed (5 – 20 minutes). RNA was extracted by mixing the homogenate with 1:1 volume of chloroform/isoamylalcohol (24:1). Supernatant was collected after centrifugation at 10,000 *g* for 15 minutes at 4°C. Supernatant was collected after centrifugation 10,000 *g* for 15 minutes at 4°C and the chloroform/isoamylalcohol extraction step was repeated. RNA was precipitated with 3 M LiCl and 1 % (v/v) beta-mercaptoethanol at −20 °C overnight. Samples were centrifuged for one hour at 4 °C at 20,000 *g*. Pellets were washed with ice cold 70 % ethanol and then dissolved in RNAse-free water. DNase treatment was performed using a TURBO DNase Kit according to manufacturer instructions (Thermo Fisher, #AM1907). Libraries were constructed with the NEBNext® Ultra™ II Directional RNA Library Prep Kit (New England Biolabs, #E7760S) and sequenced on an Illumina NextSeq2000 platform to generate a minimum of 25-30 million 150-bp paired-end reads per sample.

The RNA-seq samples were processed according to the methods described by (Perroud et al., 2018) and updated (Haas et al., 2020). The software packages used by the RNA-seq pipeline were updated to the most recent versions: gmap-gsnap (Wu and Nacu, 2010) version 2021-12-17; samtools (Li et al., 2009) version 1.15.1. Read count software was exchanged by subread featurecounts (Liao et al., 2014), version 2.0.3. Published RNA-seq datasets for WT male and female gametophytes were downloaded and processed in the same manner. Differential gene expression analysis was performed using DESeq2 version 1.30.1 (Love et al., 2014). Differentially expressed genes were classified as having a log_2_ fold-change difference of 1 and a *P* value < 0.05 after correction for multiple testing (i.e., an adjusted *P* value). We considered a conservative set of 94 sex-biased genes (SBGs) that were commonly sex-biased (i.e., differentially expressed in WT male or WT female gametophytes) in at least two out of three independent public datasets (Table S2) (). Male and female SDR genes, gametologs and PAR genes were described previously (Gueno et al., 2022). RNA-Seq heatmaps were generated using the pheatmap package in R (https://github.com/raivokolde/pheatmap). Violin plots were generated using the ggplot2 package in R (https://ggplot2.tidyverse.org) while dot plots were generated using base R.

### Small RNA analysis

WT sporophyte (Ec17) and *oro;oro* mutant (Ec581) tissue was grown and frozen as described in the RNA-seq section. 50 mg of tissue was used for total RNA isolation employing a modified version of the CTAB-based method described above. 1 mL of CTAB buffer was used for homogenization while precipitation was carried out with a 1:1 volume of iso-propanol at −20°C overnight to preserve small RNA molecules. DNase treatment was performed using a TURBO DNase Kit according to manufacturer instructions (Thermo Fisher, #AM1907). Total RNA was purified with RNA clean and concentrator columns (Zymo Reseach, #R1013) following manufacturer instructions with critical modifications to the washing steps. Columns were washed twice with 400 μL of RNA Prep buffer and four times with 700 μL of RNA Wash Buffer. Library preparation and sequencing were carried out by Novogene. Briefly, 5’ and 3’ adaptors were ligated to small RNA ends followed by first strand cDNA synthesis after hybridization with a reverse transcription primer. Double-stranded cDNA libraries were generated through PCR enrichment. Fragments containing inserts between ∼18-40 bp were size selected and purified prior to Illumina sequencing to generate 50-bp single-end reads.

Data quality control, trimming and mapping were performed with a Snakemake sRNA pipeline (https://github.com/seb-mueller/snakemake_sRNAseq). In brief, FastQC (v0.11.7) was used to assess read quality, followed by 3’ adaptor removal using cutadapt to trim Illumina universal adapters. All sequences <18 bp and >40 bp in length were filtered and the remaining sequences mapped to the *Ectocarpus sp.* 7 reference genome (https://phaeoexplorer.sb-roscoff.fr/public/organism/ectocarpus-sp7/). Mapping was performed using Bowtie version 1.2 with no mismatches allowed (Langmead and Salzberg, 2012). Both unique and multimapped sRNAs were considered for downstream analysis. The config.yaml files used for this analysis are supplied in Supplemental Data S1. sRNA quantity was normalized as count per million (CPM) and counted over gene models using the *featureCounts* tool in the R package Subread with default parameters (Liao et al., 2014). Plots and statistical tests were performed with base R and ggplot2 1.0. DESeq2 was used to calculate differential sRNA accumulation (Love et al., 2014). Correlation analysis was carried out using the *ggscatter* function (add = “reg.line”, conf.int = TRUE, cor.coef = TRUE, cor.method = “spearman”) with the R package Ggpubr (https://cran.r-project.org/web/packages/ggpubr/index.html).

### ChIP-seq analysis

ChIP-seq profiling of H3K4me3, H3K9ac, H4K20me3, and H3K79me2 were performed as described previously (Bourdareau et al., 2022). In brief, approximately 1 g of semi-dry *Ectocarpus* tissue (corresponding to around 1,000 individuals) was fixed in seawater containing 1% formaldehyde for five minutes. Cross-Formaldehyde was eliminated by washing with fresh seawater and the cross-linking quenched by incubation in 1x PBS containing 400 mM glycine for 5 minutes. Nuclei were isolated by grinding the cross-linked tissue in liquid nitrogen, resuspended in nuclear isolation buffer (0.1% Triton X-100, 125 mM sorbitol, 20 mM potassium citrate, 30 mM MgCl2, 5 mM EDTA, 5 mM beta-mercaptoethanol, 55 mM HEPES at pH 7.5, 1x cOmplete protease inhibitor cocktails (Roche)), then gently ground in a Tenbroeck Potter. The suspension was filtered through Miracloth then centrifuged at 3000g for 20 minutes to pellet the nuclei. The nuclear pellets were washed twice with fresh nuclei isolation buffer and once with nuclei isolation buffer containing no Triton X-100. The final pellet was resuspended in 750 uL nuclear lysis buffer (10 mM EDTA, 1% SDS, 50 mM Tris-HCl at pH 8, 1x cOmplete protease inhibitor cocktails (Roche)). Chromatin was fragmented by sonicating the nuclear suspension in a microTUBE AFA Fiber Snap-Cap 6×16mm using a Covaris E220 Evolution sonicator (duty 25%, peak power 75, cycles/burst 200, duration 900 s at 4°C). The sonicated suspension was then centrifuged (14,000 *g*) for 5 minutes at 4°C to remove cell debris. The supernatant containing fragmented chromatin was collected and diluted 10 times with ChIP dilution buffer (1% Triton X-10, 1.2 mM EDTA, 16.7 mM Tris-HCl pH 8, 167 mM NaCl and 1x cOmplete protease inhibitor cocktail (Roche). The chromatin solution was split among multiple DNA-Lo Bind Tubes (Eppendorf) and incubated with antibodies on a rotator at 10RPM overnight at 4°C. All histone antibodies were purchased from Cell Signalling Technology (anti-histone H3 #4620; anti-H3K4me3 #9751S; anti-H3K9ac #9649S; anti-H3K79me2 #D15E8; anti-H4K20me3 #5737S). Immunoprecipitation was performed using an equal mix of protein A and protein G Dynabeads (Thermo Scientific, #10004D and #10002D). Following immunoprecipitation and washing steps, samples were eluted in 100uL Direct Elution Buffer (0.5 % SDS, 5 mM EDTA, 10 mM Tris-HCl pH 8, 300 mM NaCl). Cross-links were reversed by incubating the samples at 65°C overnight with intermittent shaking. The samples were digested with Proteinase K (Fischer Scientific, #11826724) and RNAse A (Roche, #10109142001) prior to DNA purification using AMPure beads (Beckman Coulter, #A63882). Libraries were constructed with the NEBNext® Ultra™ II DNA Library Prep Kit (New England Biolabs, #E7645S) and sequenced on an Illumina HiSeq 3000 platform to generate a minimum of 20 million 150-bp reads per sample.

Two biological replicates of each genotype were mapped onto the *Ectocarpus sp.* 7 reference genome (https://phaeoexplorer.sb-roscoff.fr/public/organism/ectocarpus-sp7/) using the Nextflow nf-core/chipseq pipeline v1.2.2 (Harshil Patel et al., 2021). Published ChIP-Seq datasets for WT male and female gametophytes were downloaded and processed in the same manner (Gueno et al., 2022). A MultiQC report (Ewels et al., 2016) of each run with quality control metrics of each dataset, including trimming, mapping, coverage and complexity metrics, as well as the version of each tool used in the pipeline, are included in Supplemental Data S2. For data visualization and plotting, normalised log_2_ bigwig coverage files of each histone mark relative to H3 were generated using deepTools version 3.5.1 bamCompare with a bin size of 10 bp. Biological replicates were merged for downstream analysis after confirming high correlation (Fig. S2). Cross-correlation matrices of Spearman’s correlation coefficient were generated by comparing log_2_ coverage relative to H3 using deepTools version 3.5.1 multiBigwigSummary (Ramirez et al., 2014). Bigwig coverage files were visualized along the *Ectocarpus* genome using IGV version 2.16.2 ((Robinson et al., 2011)). Averaged ChIP-Seq profiles were generated using the R package EnrichedHeatmap normalizeToMatrix (Gu et al., 2018) and plotted using a custom script or with deepTools version 3.5.1 plotProfile (Ramirez et al., 2014). Differential analysis of H3K79me2 peaks between WT gametophytes and sporophytes was performed using the R package DiffBind version 3.18 (https://bioconductor.org/packages/release/bioc/html/DiffBind.html). Hierarchical clustering of H3K79me2 profiles among the different genotypes was computed with deepTools version 3.5.1 computeMatrix using log_2_ H3K79me2 coverage relative to H3.

## Data availability

Deep-sequencing data arising from this study have been deposited in the Gene Expression Omnibus under accession code PRJNA1055718. All other data supporting the findings of this study are available from the corresponding authors upon reasonable request.

## Acknowledgements

The authors thank Remy Luthringer and Dorothee Koch for help with algal cultures. This study was funded by the Max Planck Society, European Research Council grant 864038 (SMC) and the Moore Foundation (SMC). JV thanks the International Max Planck Research School ‘From Molecules to Organisms’ for support.

## Contributions

JV: Investigation (lead); Formal analysis (supporting); Visualization (supporting).

CM: Investigation (supporting); Formal analysis (supporting); Methodology (supporting); Visualization (supporting); Writing – review and editing (supporting).

OG: Investigation (supporting).

FBH: Data curation (equal); Formal analysis (supporting).

MB: Data curation (equal); Formal analysis (lead); Methodology (equal); Supervision (equal); Visualization (lead); Writing – original draft (equal); Writing – review and editing (equal).

SMC: Conceptualization (lead); Funding acquisition (lead); Methodology (equal); Project administration (lead); Supervision (equal); Visualization (supporting); Writing – original draft (equal); Writing – review and editing (equal).

## Supplemental Figures

**Figure S1.**
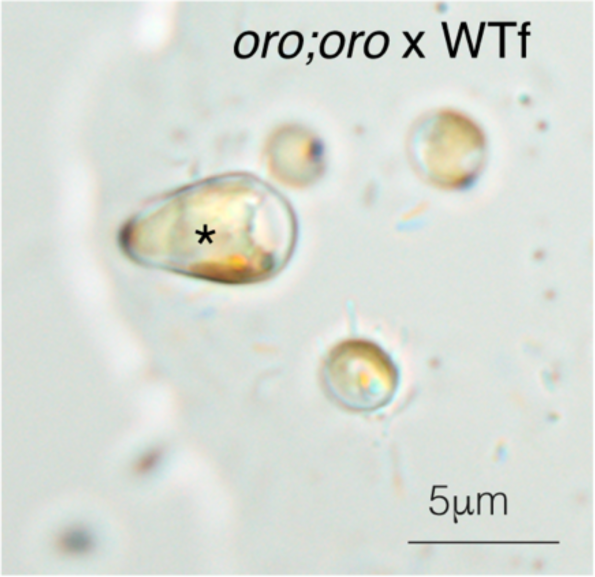
*oro;oro* mutants function as fertile male gametophytes. Image of a developing zygote (asterisk) obtained from a cross between a wild type (WT) female and diploid *oro;oro* mutant strain.

**Figure S2.**
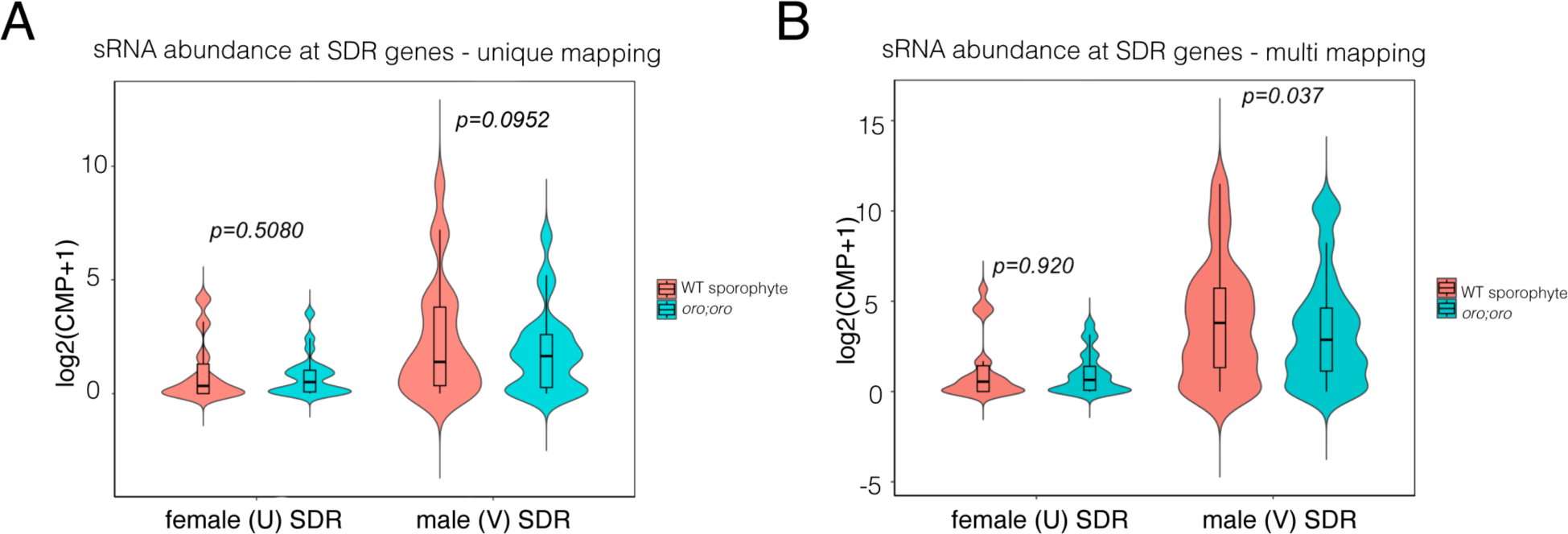
Small RNA accumulation over genes involved in sex determination. Violin plots summarising the aduncance of uniquely mapped (A) or multi-mapped (B) sRNAs on SDR genes and uniquely mapped (C) or multi-mapped (D) sRNAs on sex-biased genes. U = genes on the SDR of the female U sex chromosome, V = genes on the SDR of the male V sex chromosome, FBG = Female-biased genes, MBG = Male-biased genes. Abundance in the violin plots represents the log_2_ of the mean sRNA-seq CPM+1 values. *P* values were computed using a paired Wilcoxon signed-rank test.

**Figure S3.**
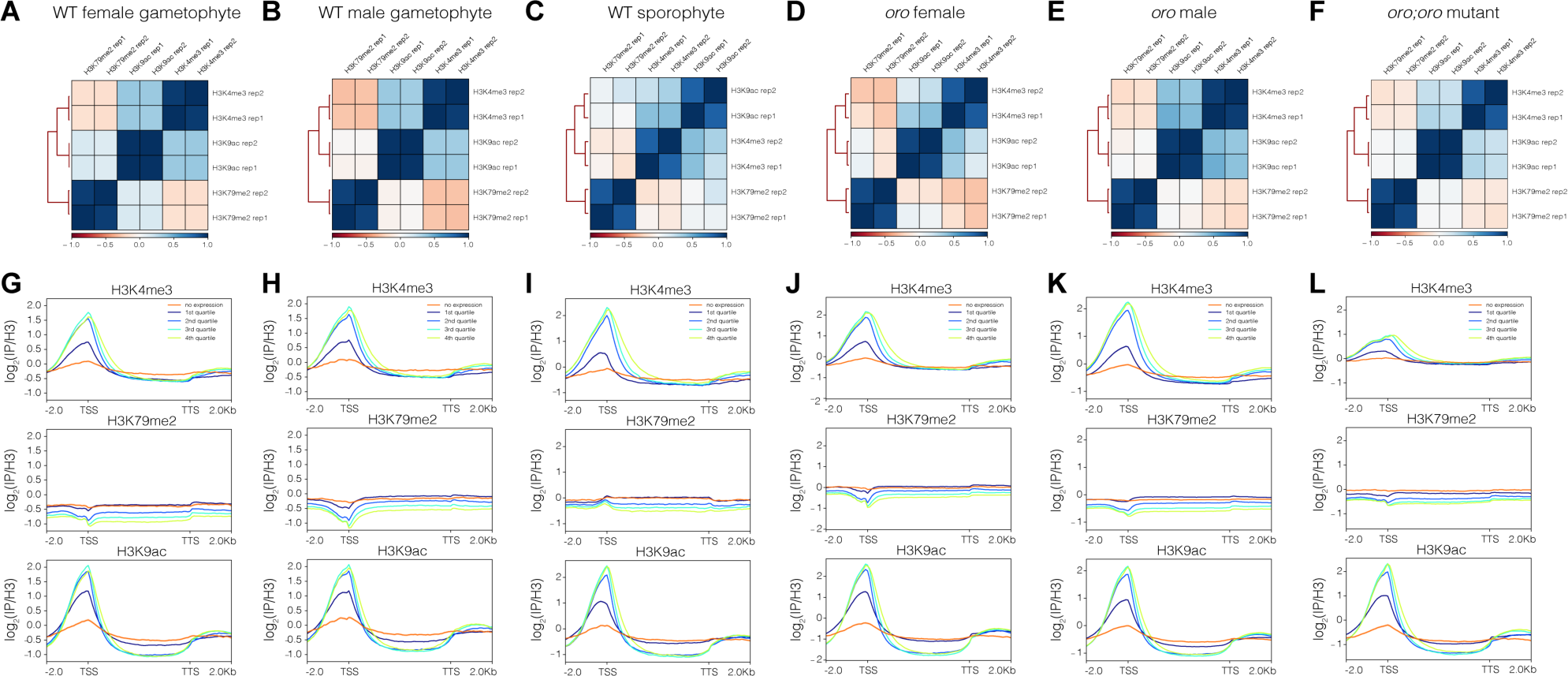
Quality control of the ChIP-seq datasets generated and/or analysed in this study. Pearson correlation matrices of the ChIP-seq replicates (A-F) and the ChIP-seq signal of H3K4me3, H3K9ac and H3K79me2 over genes sorted by the expression level in each corresponding genotype (G-L). (A,G) WT female gametophyte, (B,H) WT male gametophyte, (C,I) WT sporophyte, (D,J) *oro* females, (E,K) *oro* males and (F,L) diploid *oro;oro* mutants.

## Supplemental Tables

Table S1 – TPM values of RNA-seq data used in this study alongside differential expression data of comparisons between WT and oro gametophytes.

Table S2 – TPM values of the sex-biased genes used in this study.

Table S3 – Differentially-expressed transcripts and associated sRNAs at SDR genes.

Table S4 – H3K79me2 peaks with significant differential enrichment between WT sporophyte and WT female gametophytes.

Table S5 – H3K79me2 peaks with significant differential enrichment between WT sporophyte and WT male gametophytes.

## Supplemental Datasets

Supplemental Data S1 – Configuration files used for the processing of sRNA-seq data. Supplemental Data S2 – MultiQC reports of processed ChIP-seq data.

